# A pangenome analysis of ESKAPE bacteriophages: the underrepresentation may impact machine learning models

**DOI:** 10.1101/2024.02.19.581114

**Authors:** Jeesu Lee, Branden Hunter, Hyunjin Shim

**Affiliations:** Center for Biosystems and Biotech Data Science, Ghent University Global Campus, Incheon 21985, South Korea; Department of Biology, California State University, Fresno, 5241 N Maple Ave, Fresno, CA 93740, USA

**Keywords:** Bacteriophages, ESKAPE pathogens, Pangenome, Protein prediction, Unbalanced datasets

## Abstract

Bacteriophages are the most prevalent biological entities in the biosphere. However, limitations in both medical relevance and sequencing technologies have led to a systematic underestimation of the genetic diversity within phages. This underrepresentation not only creates a significant gap in our understanding of phage roles across diverse biosystems but also introduces biases in computational models reliant on these data for training and testing. In this study, we focused on publicly available genomes of bacteriophages infecting high-priority ESKAPE pathogens to show the extent and impact of this underrepresentation. First, we demonstrate a stark underrepresentation of ESKAPE phage genomes within the public genome and protein databases. Next, a pangenome analysis of these ESKAPE phages reveals extensive sharing of core genes among phages infecting the same host. Furthermore, genome analyses and clustering highlight close nucleotide-level relationships among the ESKAPE phages, raising concerns about the limited diversity within current public databases. Lastly, we uncover a scarcity of unique lytic phages and phage proteins with antimicrobial activities against ESKAPE pathogens. This comprehensive analysis of the ESKAPE phages underscores the severity of underrepresentation and its potential implications. This lack of diversity in phage genomes may restrict the resurgence of phage therapy and cause biased outcomes in data-driven computational models due to incomplete and unbalanced biological datasets.

## Background

Bacteriophages (phages), comprising the most abundant biological entities in the biosphere, play a crucial role in various ecological and microbial systems [1,2]. They contribute significantly to the dynamics of microbial ecosystems, influencing bacterial populations and diversity. The interconnectedness of bacteriophages with bacterial communities underscores their importance in shaping microbial dynamics, with potential consequences for human health, agriculture, and environmental processes. Despite their ubiquity, the comprehensive understanding of their genetic repertoire has been relatively limited compared to that of other organisms [2]. This systematic understudy has resulted in a significant gap in our biological knowledge, particularly regarding the multifaceted roles phages play in diverse biosystems [3,4].

While historically perceived to have limited medical relevance compared to bacteria, the significance of phages in the medical domain is experiencing a resurgence. This renewed interest is closely linked to the escalating threat of antimicrobial resistance [5], which has prompted a critical reassessment of alternative therapeutic strategies, notably the effectiveness of phage therapy. In the face of increasing bacterial resistance to traditional antibiotics, phages - viruses that infect and replicate within bacteria - have emerged as promising candidates for combating bacterial infections [6,7]. The specificity of phages in targeting particular bacterial strains, coupled with their ability to co-evolve with bacteria, presents a dynamic and potentially effective approach to counteract the challenges posed by antimicrobial resistance [8]. This shift in perspective underscores the evolving landscape of medical research, emphasizing the importance of harnessing the unique attributes of phages in addressing the pressing global concern of antimicrobial resistance.

The underrepresentation of phages across various datasets not only impedes our capacity to unravel the complex dynamics of microbial interactions in nature but also introduces biases in diverse biological models. A majority of these models, reliant on current genomic data, may inadvertently incorporate a skewed perspective, thereby limiting their accuracy and applicability. This biased outcome is analogous to the recent issues in computer vision due to dataset imbalance and bias [9,10]. For example, deep learning models have made significant strides in solving the long-standing protein folding problem in biology [11]. However, these models rely on experimental protein structures for training and testing, which are used in combination with genomic data for multiple sequence alignment [12]. If genomic databases and protein structure repositories exhibit biases toward certain organisms, computational models derived from these datasets may fail to accurately represent the biological landscape. Bacteriophages, renowned for their rich diversity of small proteins [3], represent a facet of the protein landscape that remains relatively unexplored within the current biological context (Figure 1a). Therefore, addressing this knowledge gap becomes imperative not only for advancing our understanding of phage biology but also for refining computational models essential for numerous scientific applications.

**Figure 1.**
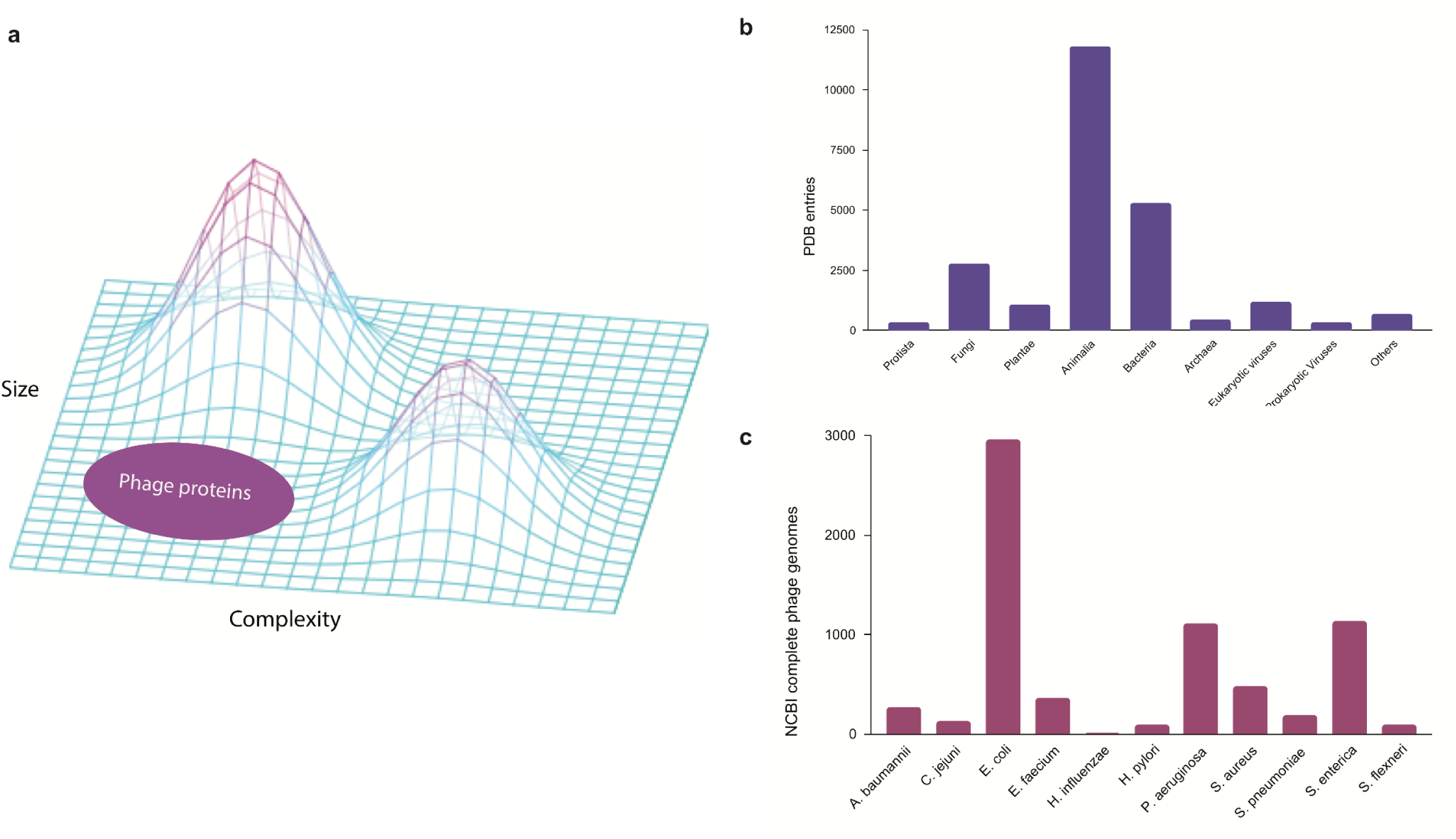
Underrepresentation of bacteriophages in the public databases. (a) Three-dimensional protein structure landscape represented by the variations in size and complexity. (b) Experimental protein structure entries by natural source organisms in the Protein Data Bank (PDB) classified at the Kingdom level. (c) Complete genomes of ESKAPE phages in the NCBI Virus database.

In this study, we examine the extent of the underrepresentation of phages in the genome and protein databases, focusing on the phages that are infecting the high-priority pathogens defined by the WHO [8]. These phages may become medically relevant as an alternative antimicrobial therapy against ESKAPE pathogens, which is an acronym for the high-priority pathogens, made up of *Enterococcus* spp., *Staphylococcus aureus*, *Klebsiella pneumoniae*, *Acinetobacter baumannii*, *Pseudomonas aeruginosa*, and *Enterobacter* spp. [13]. In this study, we included 11 bacterial species from the WHO priority list and studied the phages of these ESKAPE pathogens using genome and pangenome analyses. We first analyzed the extent of biased datasets against bacteriophages in the Protein Data Bank [14] (Figure 1b), before downloading all the complete genomes of the ESKAPE phages in the NCBI Virus database for the downstream analysis (Figure 1c).

From this study, we aim to provide a quantitative analysis in understanding the extent of underrepresented datasets of phages and engage the scientific community to focus more attention on phage-related data collection given the wide implications on diverse fields from phage therapy to machine learning models. As research advances, the multifaceted roles of bacteriophages in modulating bacterial behavior, participating in microbial community dynamics, and offering therapeutic solutions continue to unfold. This evolving understanding challenges the historical notion of limited medical relevance and positions bacteriophages as integral components of the complex microbial world with far-reaching implications for diverse fields, including medicine, ecology, and biotechnology.

## Materials and methods

### Curation of ESKAPE phage datasets

We first analyzed the underrepresentation of bacteriophages in the genome and protein databases. For the protein database, we downloaded the data distribution provided by the Protein Data Bank (PDB) [14] on the entries of experimental protein structures by natural source organisms (downloaded 2024/01/24). The definition of natural source organisms for these structures is from a natural and non-modified source. To categorize each entry to a natural source organism at the kingdom level (Figure 1b), we used a combination of human expertise and a generative AI based on the large language model [15].

Next, we curated the ESKAPE phage genome dataset by downloading the reference genomes of bacteriophages from the NCBI Virus database. Here, we define the ESKAPE phages as bacteriophages that infect the WHO priority pathogens (*Helicobacter pylori*, *Campylobacter jejuni*, *Salmonella enterica*, *Streptococcus pneumoniae*, *Haemophilus influenzae*, *Shigella flexneri*) encompassing the ESKAPE (*Enterococcus* spp., *Staphylococcus aureus*, *Klebsiella pneumoniae*, *Acinetobacter baumannii*, *Pseudomonas aeruginosa*, and *Enterobacter* spp.) [16]. Some pathogens, such as *Klebsiella pneumoniae* and *Neisseria gonorrhoeae*, were omitted from the list as they do not have associated bacteriophages in the NCBI virus database. All the RefSeq genomes with the specified host were downloaded for each pathogen species as whole genome sequences and protein sequences (downloaded 2022/10/18).

The genomes of each ESKAPE phage were analyzed using several biological features and statistical measures (Table S1). The biological features included phage ID, phage type, and DNA type, and the statistical measures included GC content, AT content, GC/AT ratio, number of proteins, and sequence length. These measures were summarized into the most common phage type, the most common DNA type, GC content, and the number of open-reading frames (ORF) for each host ESKAPE pathogen (Table 1).

**Table 1.**
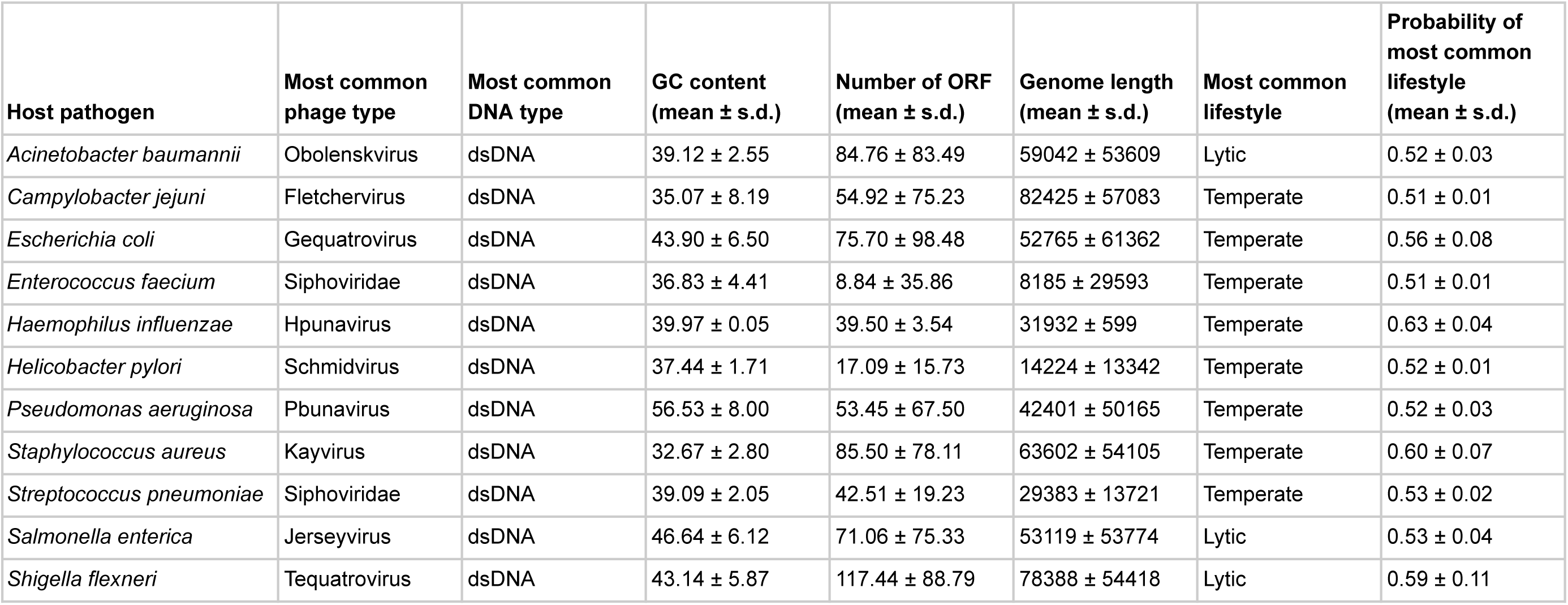
Summary statistics of phage genomes associated with the ESKAPE pathogens.

| Host pathogen | Most common phage type | Most common DNA type | GC content (mean ± s.d.) | Number of ORF (mean ± s.d.) | Genome length (mean ± s.d.) | Most common lifestyle | Probability of most common lifestyle (mean ± s.d.) |
| --- | --- | --- | --- | --- | --- | --- | --- |
| *Acinetobacter baumannii* | Obolenskivirus | dsDNA | 39.12 ± 2.55 | 84.76 ± 83.49 | 59042 ± 53609 | Lytic | 0.52 ± 0.03 |
| *Campylobacter jejuni* | Fletchervirus | dsDNA | 35.07 ± 8.19 | 54.92 ± 75.23 | 82425 ± 57083 | Temperate | 0.51 ± 0.01 |
| *Escherichia coli* | Gequatrovirus | dsDNA | 43.90 ± 6.50 | 75.70 ± 98.48 | 52765 ± 61362 | Temperate | 0.56 ± 0.08 |
| *Enterococcus faecium* | Siphoviridae | dsDNA | 36.83 ± 4.41 | 8.84 ± 35.86 | 8185 ± 29593 | Temperate | 0.51 ± 0.01 |
| *Haemophilus influenzae* | Hpunavirus | dsDNA | 39.97 ± 0.05 | 39.50 ± 3.54 | 31932 ± 599 | Temperate | 0.63 ± 0.04 |
| *Helicobacter pylori* | Schmidvirus | dsDNA | 37.44 ± 1.71 | 17.09 ± 15.73 | 14224 ± 13342 | Temperate | 0.52 ± 0.01 |
| *Pseudomonas aeruginosa* | Pbunavirus | dsDNA | 56.53 ± 8.00 | 53.45 ± 67.50 | 42401 ± 50165 | Temperate | 0.52 ± 0.03 |
| *Staphylococcus aureus* | Kayvirus | dsDNA | 32.67 ± 2.80 | 85.50 ± 78.11 | 63602 ± 54105 | Temperate | 0.60 ± 0.07 |
| *Streptococcus pneumoniae* | Siphoviridae | dsDNA | 39.09 ± 2.05 | 42.51 ± 19.23 | 29383 ± 13721 | Temperate | 0.53 ± 0.02 |
| *Salmonella enterica* | Jerseyvirus | dsDNA | 46.64 ± 6.12 | 71.06 ± 75.33 | 53119 ± 53774 | Lytic | 0.53 ± 0.04 |
| *Shigella flexneri* | Tequatrovirus | dsDNA | 43.14 ± 5.87 | 117.44 ± 88.79 | 78388 ± 54418 | Lytic | 0.59 ± 0.11 |

To visualize the statistical information, we created boxplots for two variables: phage genome lengths and the number of open reading frames (Figure 2a,b). Additionally, we created scatter plots to illustrate the relationship between these two variables (Figure 2c,d). Each data point on the plot represents every ‘phage ID’, with the phage genome lengths plotted along the x-axis and the number of open reading frames plotted along the y-axis. The scatter plots included data from all 11 species, where each species was displayed in a different color. The statistics from the *E. coli* phages were plotted separately due to the high number of data points (Figure 2d). Then, we used a ‘LinearRegression’ class to fit a model that represented the linear relationship. By using this model, we calculated the regression equation and the R^2^ value. The scatter plot with an overlayed regression line helped to identify whether the variables have a positive, negative, or no apparent relationship.

**Figure 2.**
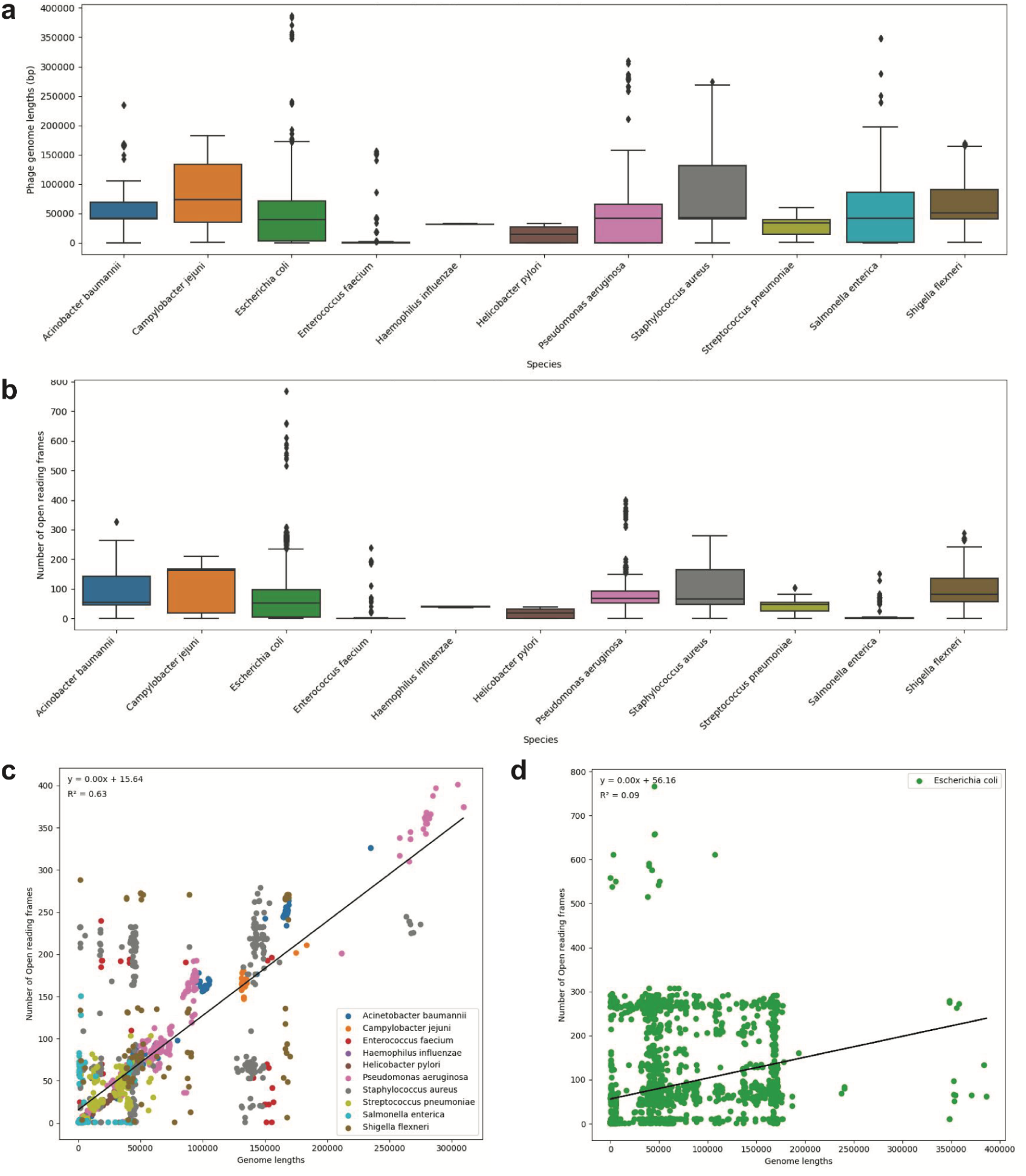
(a) Boxplot of phage genome lengths (bp) of phages by their host ESKAPE pathogen. (b) Boxplot of open reading frames of phages by their host ESKAPE pathogen. (c) Scatter plot of phage genome lengths (bp) versus open reading frames, colored by each host ESKAPE pathogen with an overlayed regression line. (d) Scatter plot of *E. coli* phage genome lengths (bp) versus open reading frames with an overlayed regression line.

### Gene function analysis of ESKAPE phages

To observe the function of diverse genes in the ESKAPE phage genomes, the fasta files were processed with Biopython to extract gene information such as gene ID, virus ID, gene name, virus name, virus genus, and family. We extracted the gene keywords such as ‘DNA’ or ‘polymerase’ from the gene names. A vast majority of the gene names contain the keyword ‘hypothetical’. A hypothetical gene is a predicted gene that is likely to be expressed in organisms, but which has never been characterized for biochemical function [17]. Therefore, keywords related to hypothetical genes were excluded from the subsequent processing.

To visualize these gene keywords and names, we generated a keyword heatmap and a horizontal bar chart for each ESKAPE gene genome, respectively. The genes with the keyword ‘putative’ also have unknown functions like hypothetical genes, but they have similar properties to already existing genes [18]. Since they are assumed to be functional genes, this keyword was kept for further study. The visualization of these genes was done in three sets; the first set includes only functional genes excluding hypothetical and putative genes (Figure S1), the second set includes only putative genes (Figure S2), and the third set includes all functional and putative genes (Figure S3).

Subsequently, we summarized the gene functions into a heatmap to compare the relative magnitudes of genes co-occurring in the ESKAPE phage genomes (Figure 3). Gene keywords from each ESKAPE gene genome were downloaded as CSV files, which were processed into ‘word’ and ‘value’ columns. To calculate the frequency of common words within 22 CSV files (11 CSV files were downloaded from each set: all genes and only functional genes), we used the ‘Counter’ object. Each word was treated as a unique key and its occurrences are computed individually, for example, ‘holin’ or ‘putative holin’. We created a list of regular expression patterns by selecting the 32 words with the highest frequencies from the extracted unique keys. Each target word in the list was processed using ‘\b{}\b’ to indicate a word boundary in the regular expression pattern. These patterns were then used to search for words in the previous CSV files that fully matched the pattern. Finally, we merged all the data into a single data frame and generated the heatmap, which provided us with a visual pattern to easily grasp the relationships between the data.

**Figure 3.**
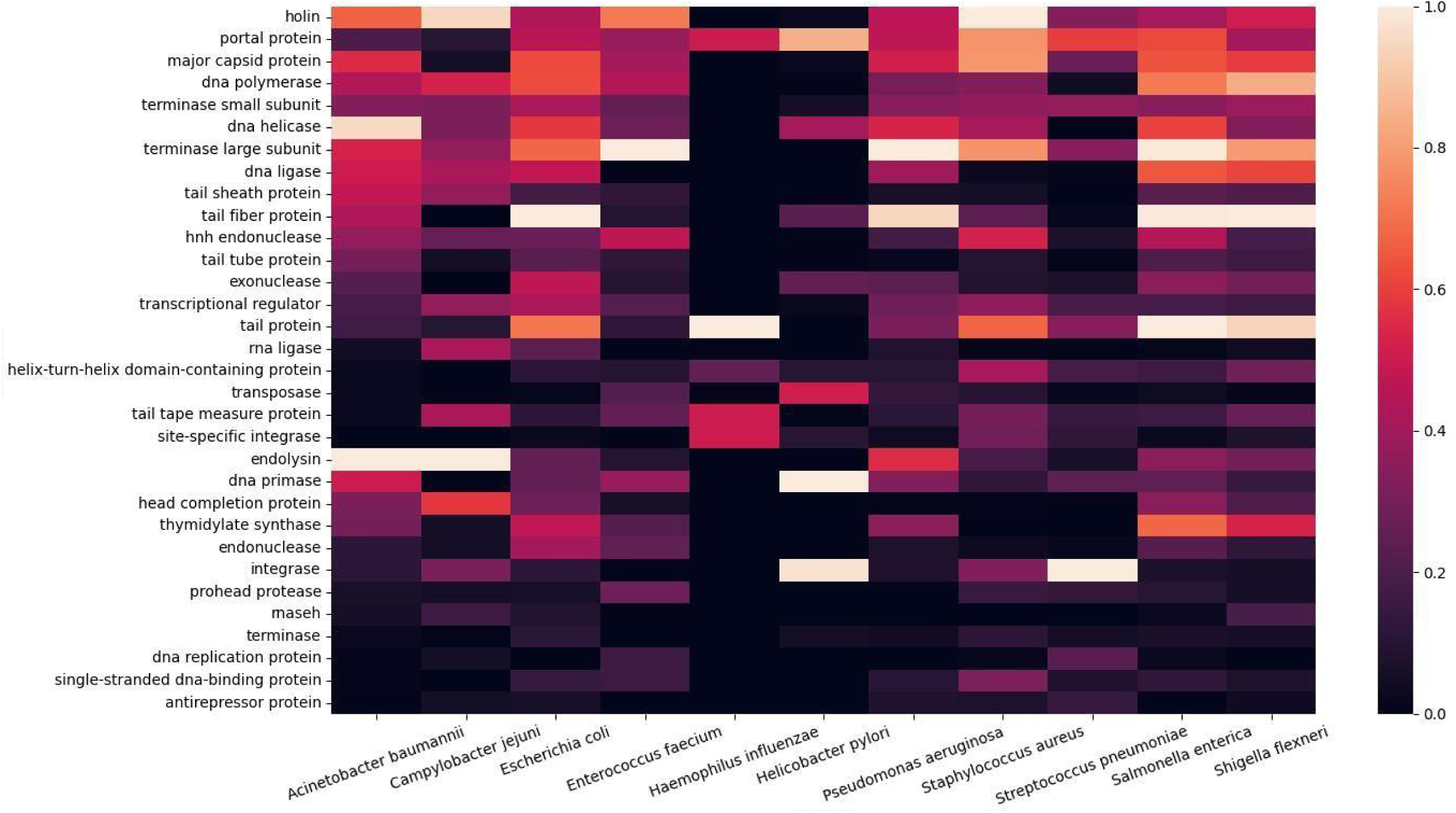
Heatmap of co-occurring functional genes from the ESKAPE phage genomes.

### Genome visualization and phage lifestyle prediction

We used a genome viewer to visualize the genomic architecture of each phage genome. In this process, the untranslated reference sequence genome fasta file and translated reference sequence protein fasta file were separated into every single IDs of phages. The translated reference sequence protein fasta file contains the information about IDs of phage and locations where the protein of each phage is translated. To simplify reference sequence protein fasta files, phage IDs were extracted from the untranslated reference sequence genome fasta files through Python. Using the phage IDs in the list, new files containing all translation sites per phage ID were created. With the DNA feature viewer algorithm, linear graphs were made with whole genome data while omitting hypothetical proteins (Figure S4).

PHACTS is a computational approach that classifies the lifestyle of bacteriophages [19]. The lifestyle of a phage can be classified as virulent (lytic) or temperate (lysogenic). For the virulent and temperate types, a phage genome is annotated as either ‘Lytic’ or ‘Lysogenic’, respectively, with a computed probability. PHACTS predicts the lifestyle of a bacteriophage based on the genome content through two training sets and the Random Forest method [20,21]. One of the training sets of the query protein sequences was selected from the newly edited fasta files. The other training set is based on the known phage sequence data from PHANTOME [22], providing a complete phage genome with two training sets. The Random Forest algorithm of PHACTS classifies the lifestyle of phages by creating multiple decision trees. Then, the known phage sequence data is loaded into decision trees through the bootstrapping method, which randomly picks data by resampling. Randomly selected query protein sequences from the newly edited fasta files are matched with the known phage sequence data to complete the decision trees. Finally, the tree with the highest computed probability is chosen as the final prediction model. To keep the accuracy and stable runtime, replicate iterations are executed 10 times. Among 10 iterative replicates, the sole consensus result values are determined as the confident data [19].

### Pangenome analysis of ESKAPE phage genomes

We used IPGA (v1.09) to analyze, compare, and visualize the pangenome of ESKAPE phages [23]. This web tool features a scoring system that evaluates the reliability of profiles generated by different pangenome methods. We ran several pangenome packages with the phages of *Salmonella enterica* as an initial trial, but the only package that worked on phage genomes as input was PEPPAN [24]. Thus, all the other phage genomes of the ESKAPE pathogens were analyzed using PEPPAN as the pangenome analysis option (Figures 4 and S5). As pangenome analysis requires four or more individual genomes, the phages of *H. influenzae* with only two individual genomes were excluded from the pangenome analysis. In addition, IPGA also implements several downstream comparative analysis modules and genome analysis modules, including Average Nucleotide Identity (ANI) that measure nucleotide-level genomic similarity between the coding regions of genomes [25].

**Figure 4.**
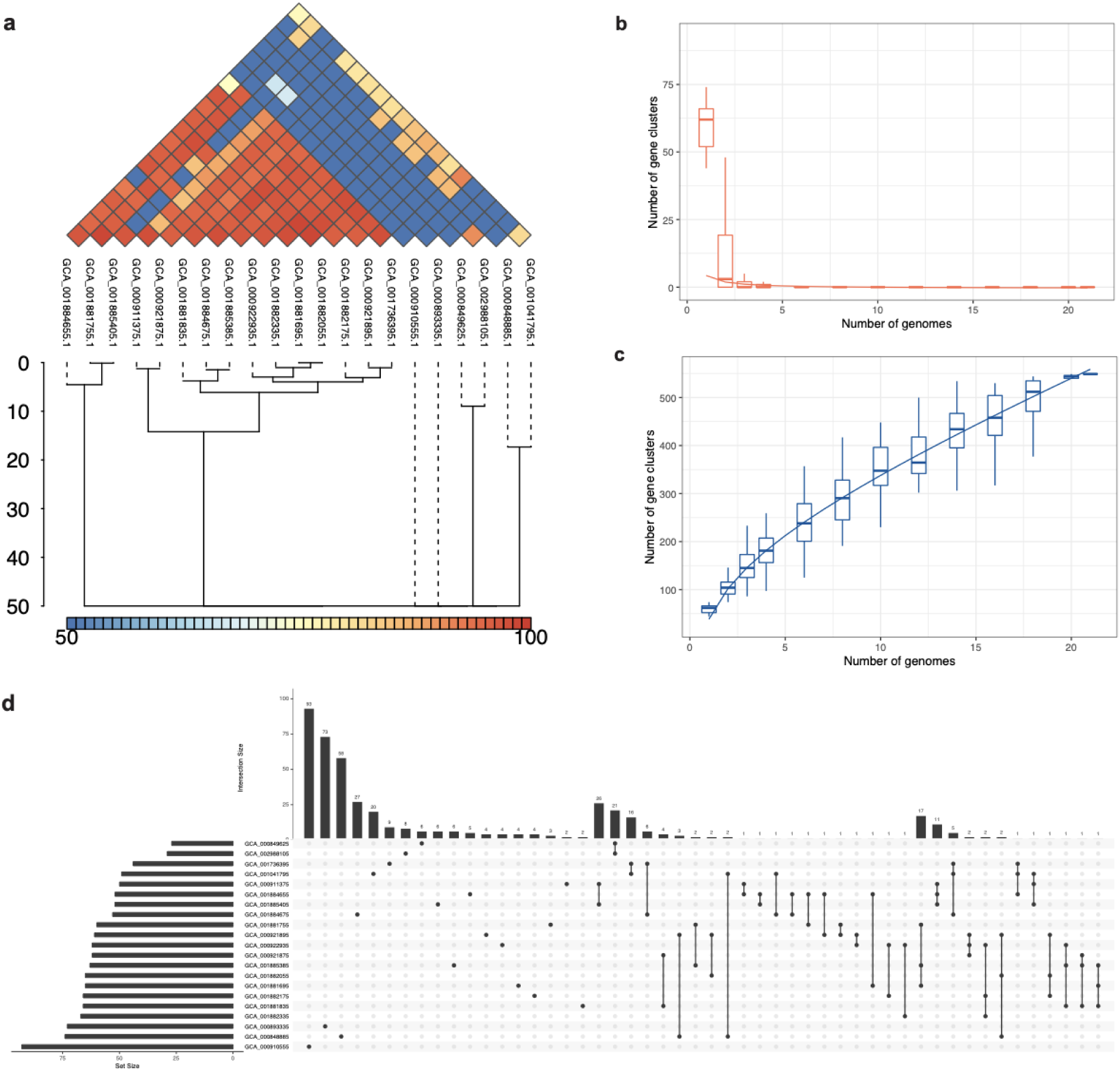
Pangenome analysis of the ESKAPE pathogens. (a) Average Nucleotide Identity (ANI) in *S. pneumoniae* phages. (b) Core genome clusters in *S. pneumoniae* phages. (c) Pangenome clusters in *S. pneumoniae* phages. (d) Set sizes and intersection sizes in *S. pneumoniae* phages.

MMseqs2 is a deep learning-based software to search and cluster huge sequence sets, with a highly efficient clustering module to group similar sequences into clusters [26]. For clustering whole-genome sequences of these ESKAPE phages, we used a clustering module of MMseqs2 that is highly efficient at grouping similar sequences into clusters. It employs an iterative clustering approach, progressively merging sequences into clusters while optimizing a predefined objective function to achieve accurate and scalable clustering results. We created a database containing the genomes of phages against each ESKAPE pathogen and clustered these genomes with a minimum sequence identity of 70%.

### Clustering of phage protein sequences by similarity

For clustering protein sequences, a clustering in linear time called the Linclust algorithm in MMseqs2 was used for fast clustering, which reduces the time complexity O(NK) to O(N), where K is the parameter that indicates the final number of clusters. The Linclust algorithm operates by generating a table that consists of the k-mer, the sequence identifier, and the sequence position, and sorting the table by k-mer to identify groups of sequencing sharing the same k-mer in quasi-linear time. Then, the longest sequence is selected as the center sequence among the sequences that share the same k-mer. The groups are merged around the center sequence and the sequence groups are compared by the global Hamming distance and gapped local sequence alignment. Finally, the representative sequence and aligned sequences are determined through the incremental greedy algorithm.

We used two settings of sequence identity thresholds (50% and 90%, respectively) for clustering as shown below. For the sequence identity of 50%, the mode was set as 1 which enables the alignment to cover at least 50% between query and target.

mmseqs easy-cluster examples/DB.fasta clusterRes tmp --min-seq-id 0.5 -c 0.5 --cov-mode 1

For the sequence identity of 90%, the mode was set as 0 which enables the alignment to cover at least 80% between query and target.

mmseqs easy-cluster examples/DB.fasta clusterRes tmp --min-seq-id 0.9 -c 0.8 --cov-mode 0

We compared the results from these two sets with the different sequence identity thresholds. We determined that the sequence identity threshold of 90% was too stringent, thus the following structure analysis was conducted with the set with the sequence identity threshold of 50%. The representative sequences were extracted from the resulting MMseqs2 file for AlphaFold structure prediction.

### AlphaFold-predicted structures of representative inhibitor phage proteins

AlphaFold is a deep learning-based program that could predict three-dimensional structures of proteins [12]. We used AlphaFold to gain insight into the structure of representative inhibitor phage proteins (Figures 5 and S6). We performed AlphaFold using Google Colab and generated protein structure predictions from the representative protein sequences. To identify only the inhibitor proteins of lytic phages, we selectively screened lytic phages of the ESKAPE pathogens (Table S2) and identified the ‘Phage ID’ of the lytic phages that corresponded to the ‘Protein ID’ of the inhibitor phage proteins (Table S3).

**Figure 5.**
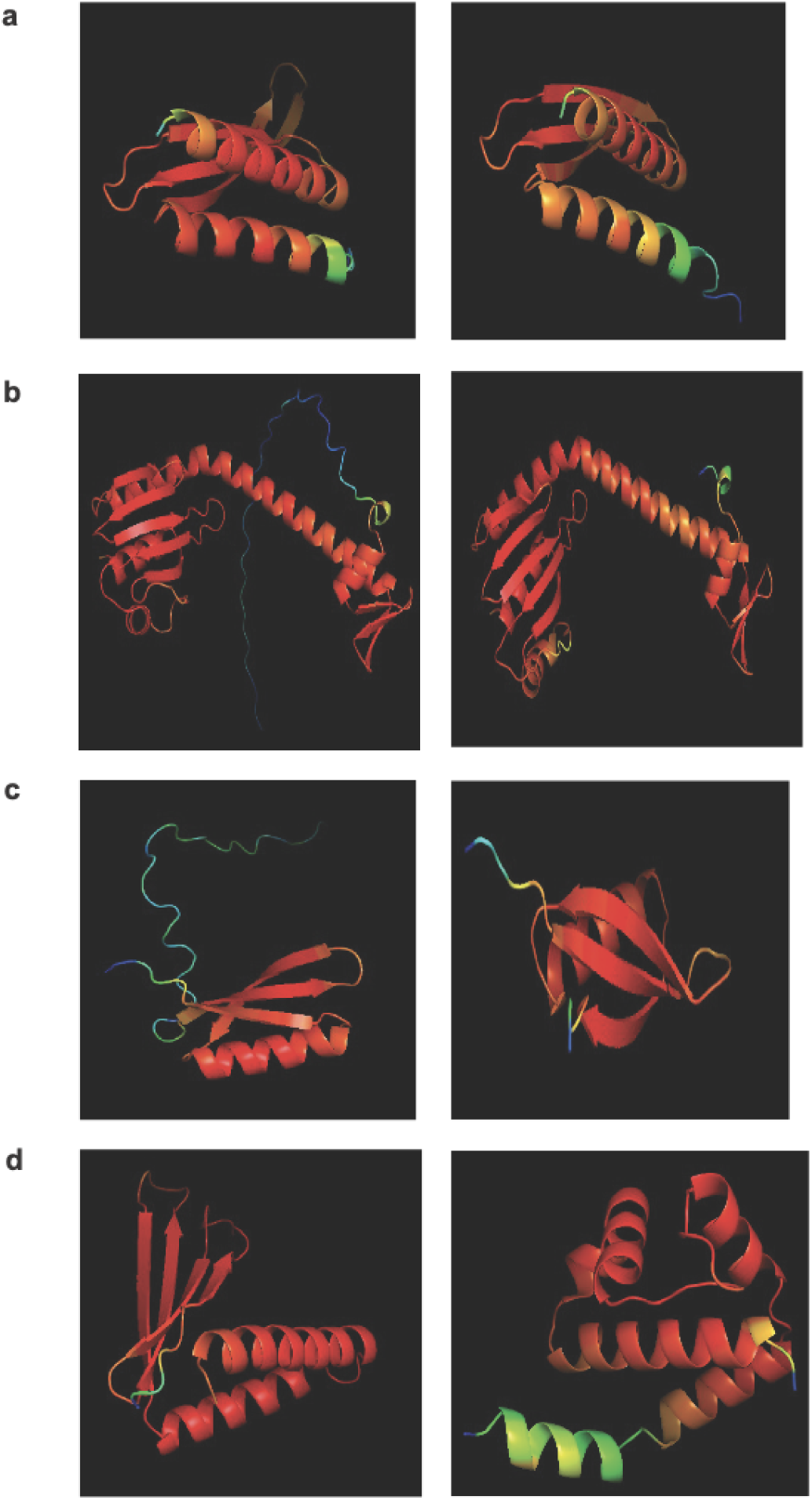
AlphaFold-predicted inhibitor proteins from lytic phages of the ESKAPE pathogens. (a) Inhibitors of host energy from *S. enterica* phages. (b) Inhibitors of host transcription from *E. coli* phages and *S. enterica* phages. (c) Inhibitors of host translation from *S. enterica* phages. (d) Inhibitor of host toxin/antitoxin system from *S. enterica* and anti-sigma factor from *E. coli* phages.

Subsequently, the PDB files of the predicted protein structures were downloaded, and we observed the three-dimensional representation of protein structures with the Pymol, including alpha helices, beta sheets, and tertiary structures. In addition, we analyzed the temperature factor column in the PDB file describing the per-residue Local Distance Difference Test (pLDDT) of each residue [12]. These scores are represented on a spectrum, with higher certainty depicted in red and lower certainty shown in blue.

## Results

### Bacteriophages are highly underrepresented in the databases

In the Protein Data Bank (PDB), we discovered that the experimental structures of bacteriophage proteins are highly underrepresented when classified at the kingdom level (Figure 1b). The bar chart of the PDB entries by natural source organism shows that the kingdom of Animalia is highly represented, particularly biological model organisms such as *Homo sapiens* (3364 entries) and *Mus musculus* (1265 entries). Furthermore, the kingdoms that contain pathogens against humans such as Bacteria and Eukaryotic viruses are well represented in the PDB database with entries of 5262 and 1165, respectively. The least represented kingdoms are Protista, Archaea, and Prokaryotic viruses (i.e. bacteriophages) with entries of 294, 415, and 309, respectively. For the ESKAPE phages, we found only few to no entries for *Acinetobacter baumannii* (0), *Campylobacter jejuni* (0), *Escherichia coli* (73), *Enterococcus faecium* (0), *Haemophilus influenzae* (0), *Helicobacter pylori* (4), *Pseudomonas aeruginosa* (10), *Salmonella enterica* (12), *Shigella flexneri* (4), *Staphylococcus aureus* (8), *Streptococcus pneumoniae* (3).

For the genome database, this study reveals a paucity of complete phage genomes associated with the WHO priority list pathogens in the NCBI Virus database (Figure 1c). For example, *H. influenzae*, a gram-negative bacterium causing pneumonia, meningitis, or bloodstream infections [27], has only two known complete phage genomes. Furthermore, these phages exhibit temperate lifestyles (Table 2), limiting their suitability for phage therapy applications. Despite the evident advantages of phage therapy, several challenges persist, especially in the realm of fundamental research. A notable challenge is the inadequate comprehension of the diversity of bacteriophages within their natural habitats, such as human microbiomes. This scarcity of diverse phage genomes poses a substantial impediment to the development of effective phage therapy treatments tailored to specific bacterial infections.

**Table 2.**
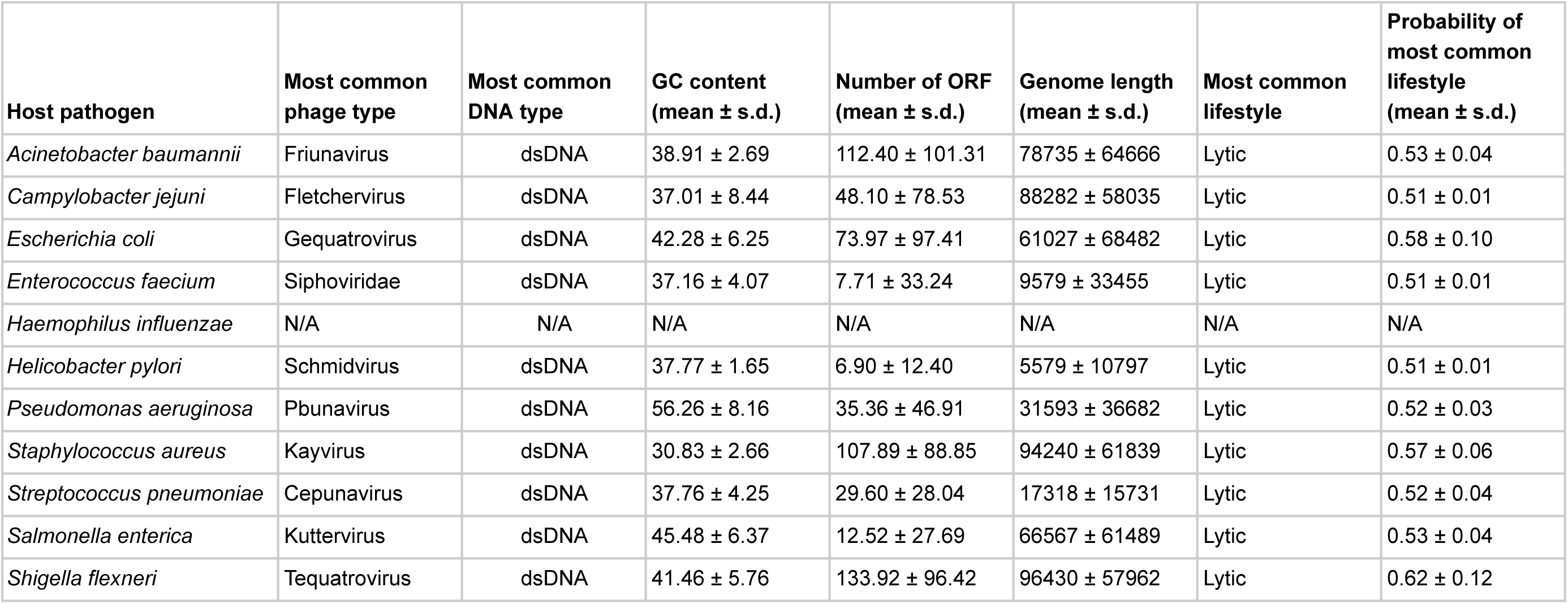
Summary statistics of lytic phage genomes associated with the ESKAPE pathogens.

Boxplots provided us with a concise overview of statistics (distribution, median, quartiles, and presence of outliers), allowing for easy comparison and interpretation (Figure 2). The boxplots of the phage genome lengths by the host show that most phages have genome lengths below 50 kbp (Figure 2a). Notably, the phages of *E. coli*, *P. aeruginosa*, and *S. enterica* have high variations in the genome length, with some phages reaching above 200 kbp in length [28]. The boxplots of the number of ORFs by the host show that most phages have protein lengths below 100 bp (Figure 2b). Notably, the phages of *E. coli* and *P. aeruginosa* have high variations in the protein length, with some proteins reaching above 400 bp in length. This result shows that most ESKAPE phages have small genome lengths packed with small proteins [29,30]. The scatter plots with an overlayed regression line show that the relationship between the genome length and the number of ORFs is positive, but this relationship only has a coefficient of determination, or R^2^, value of 0.63 for the ESKAPE phages, except that of a much lower value of 0.09 for *E. coli* phages as shown in Figure 2c and Figure 2d, respectively. This R^2^ value measures the goodness of fit of this regression model, and it shows that only 63% of the variance in the number of ORFs can be explained by the genome length variables in the ESKAPE phages, except in the *E. coli* phages with a much lower value at 7%. The variation of the number of ORFs versus the genome length appears to be much higher in the *E. coli* phages, with only many small phages having a larger number of proteins than expected by the regression model, and vice versa (Figure 2d).

### Some ESKAPE phages have genes with antimicrobial activities

As the first exploratory analysis, we created heatmaps visualizing the top keywords in the annotations of the ESKAPE phage genomes. We searched for keywords containing ‘inhibit’ or ‘anti’ that are potentially related to the antimicrobial activities of phages against their hosts. The ESKAPE phage with the most antimicrobial keywords such as ‘inhibit’ or ‘anti’ within the functionally validated genes was *E. coli* (Figure S1). Even after adjusting for the large dataset of available genomes, the *E. coli* phages have 10 or 20 times more genes with the antimicrobial keywords as compared to the *P. aeruginosa* phages with half the available genomes. The phages of *S. enterica* contain the next most abundant genes associated with antimicrobial activities. Notably, the phages of *H. pylori* and *C. jeuni* have almost no functional genes with antimicrobial activities, which may be explained by the lack of available phage genomes in the NCBI Virus database (Figure 1). However, the phage genomes of *C. jejuni* contain a small number of putative antimicrobial genes, including ‘anti-holin’, in the putative gene list (Figure S2).

This finding remains the same when the putative genes were included in the analysis (Figure S2). The putative genes containing the keywords ‘inhibit’ or ‘anti’ show the potential for novel antimicrobial activities from the ESKAPE phages. For example, the ‘anti-proliferative’ gene only appears in the list of putative genes, and, interestingly, this gene may be involved in the regulation of cell growth and development [31,32].

Subsequently, we analyzed the most frequent co-occurrence of functional proteins in the ESKAPE phage genomes (Figure 3). The heatmap shows the top 30 functional proteins that are shared between the ESKAPE phage genomes in this study. This heatmap shows that most annotations are related either to the replication function of bacteriophages, such as polymerase and helicase, or to the assembly function of bacteriophages, such as capsid and tail.

Finally, we analyzed the gene content of these phages further examining individual genes through bar graphs (Figure S3). The bar charts show the 30 most frequent genes in each ESKAPE phage. These bar charts reveal some genes of interest may have inhibitory functions. Several ESKAPE phages have several antimicrobial genes that may disrupt the vital functions of host bacteria. For example, *A. baumannii*, *E. coli*, *S. enterica*, and *S. flexneri* have anti-sigma factors that bind to sigma factors and inhibit transcriptional activity in regulating prokaryotic gene expression [33]. Moreover, *E. coli*, *S. enterica*, and *S. flexneri* have host polymerase inhibitors that may disrupt the transcriptional activities of host bacteria [34]. These genes are likely to be antibacterial as the disruption of transcription leads to cell cycle arrest [35]. These phages also possess other genes that are involved in counter-defensive activities against the host defense mechanisms, such as anti-restriction genes [36], host protease inhibitors [37], anti-repressor [38], and anti-crispr genes [39–42].

### Databases have a limited number of lytic ESKAPE phages

We visualized the genomic architecture of the ESKAPE phages to explore the gene content. The genome maps of these ESKAPE phages revealed that many of these phages have similar gene content, and many of these phages are nearly identical. Remarkably, these phage genomes also share taxonomic relationships, resulting in even fewer unique genomes for potential therapeutic use. For instance, our analysis indicates two phage genomes infecting *H. influenzae* are nearly identical in gene content (Figure S4).

Given the lack of antimicrobial genes in the ESKAPE phage genomes, we explored the lifestyle of these phages (Tables 2 and S2). Identifying the lifestyle of a phage traditionally relies on labor-intensive and costly culturing techniques. However, these methods are not only time-consuming but also impractical for phage genomes derived from environmental sequencing. We used a computational approach to predict the lifestyle of the ESKAPE phages [19]. Phages demonstrate two main lifestyles: virulent (lytic) and temperate (lysogenic). This computational approach leverages the gene content inherent to phage genomes to predict the lifestyle of a phage.

According to the computational prediction, there is only a limited number of lytic phages against all the ESKAPE pathogens (Table S2). For example, the phages of *C. jejuni*, *E. faecium,* and *H. pylori* only have one lytic phage in the database. Other ESKAPE pathogens such as *A. baumannii* and *S. pneumoniae* also have four and two lytic phages, respectively. More importantly, *H. influenzae* has no lytic phages that are predicted by the computational method. The medical importance of bacteriophages lies in their potential applications in phage therapy, a field that explores the use of bacteriophages to combat bacterial infections. Bacteriophages exhibit specificity in targeting bacterial strains, making them attractive candidates for precision medicine in treating bacterial infections. Their ability to infect and lyse bacteria provides a natural and tailored approach to controlling pathogenic bacteria, including antibiotic-resistant strains. The biological features of the lytic phages were summarized into the most common phage type, the most common DNA type, GC content, and the number of open-reading frames (ORF) for each host ESKAPE pathogen (Table 2).

### Pangenome analysis shows the ESKAPE phages share many core genes

For the ESKAPE phages that share the same host, we used the clustering method to automatically cluster their genomes by similarity at the nucleotide level. When we clustered these phages with the minimum sequence identity of 70%, it revealed that the number of individual phages can be reduced into a small number of clusters (Figure 4a). For example, 263 individual phages infecting *A. baumannii* can be reduced into 3 clusters with the clustering method. This shows that the NCBI virus database only contains 3 unique phages against the high-priority pathogen of *A. baumannii*. Similarly, the genome maps of phages infecting *A. baumannii* can also be divided into a few categories based on visualization (Figure S4). Notably, many phages share the same gene content and even the same gene arrangement. However, the public databases do not compute the similarity between these phages as there is no consensus on how to classify these phages into different ‘strains’.

The objective of pangenome analysis is to assess the diversity of all genes and genomic structures across genomes within a particular clade. The primary and pivotal step in this analysis involves clustering orthologous genes. These gene clusters are subsequently categorized into three groups based on their presence in the specified sets of genomes: core genes, accessory genes, and unique genes. Multiple packages or web services have been developed for pangenome analyses of eukaryotic and prokaryotic genomes [24,43–53], but we found no method that is specifically designed for phage genomes. We chose an integrated prokaryotes genome and pangenome analysis web service called IPGA that allows phage genomes as input for pangenome analysis, downstream analysis, and visualization of the target genomes [23].

The first step of IPGA was a quality control module that removes all low-quality genomes and performs a taxonomic assignment for each genome. IPGA then predicted genes of all filtered genomes and used them as the input of the pangenome analysis module (Figures 5 and S5). Using IPGA, we compared the pangenome profiles created by different methods and found that PEPPAN [24] was the only pangenome software that performed well with the ESKAPE phage genomes as input data. Not all phage genomes of the ESKAPE pathogen had pangenome analysis results, such as *H. influenzae* and *E. faecium*, due to the lack of individual genomes. The ESKAPE phages of *E. coli* and *S. pneumoniae* failed to give any meaningful results due to the large input dataset. The pangenome results show that the ESKAPE phage genomes have a high number of core genes that are shared as the intersection between the phages infecting the same host. For example, the core genome clusters in *S. pneumoniae* phages decrease rapidly (Figure 5b) while the pangenome clusters in *S. pneumoniae* phages increase slowly (Figure 5c). This relative difference in the incline and decline rates indicates that more core genes are shared between the phage genomes of this pathogen than the accessory or unique genes. The set sizes and intersection sizes also reflect the size of the core genome versus the size of the pangenome in the *S. pneumoniae* phages (Figure 5d). This trend is observed in the pangenome analysis of the other ESKAPE phages, including *C. jejuni*, *H. pylori*, and *S. flexneri* (Figure S5).

We also conducted the downstream comparative genomic analysis modules on the filtered genomes and gene clusters, including the phylogenetic analysis module, core gene allele analysis, and average nucleotide identity (ANI) calculation module. Average Nucleotide Identity (ANI) is a measure of nucleotide-level genomic similarity between the coding regions of two genomes. The ANI statistics show that many phage genomes share high genomic similarity at the nucleotide level. For example, the ANI analysis of *S. pneumoniae* phages shows that there are 6 clusters of phage genomes with genomic similarity below 70% at the nucleotide level (Figure 5a).

### Protein structure analyses show the underexplored structure landscape

After the pangenome analysis of the ESKAPE phage genomes, we clustered the functional and putative protein sequences with antimicrobial activities. Subsequently, we predicted the three-dimensional protein structure of a representative protein from each cluster. When these representative proteins were visualized, we found that these proteins share several secondary and tertiary structural components despite being in different clusters in terms of genetic sequence (Figure S6).

For example, the inhibitors of host energy from *S. enterica* phages share similar tertiary structures (Figure 5a). Furthermore, the inhibitors of host transcription from *E. coli* phages and *S. enterica* phages also share similar tertiary structures (Figure 5b). Also, the inhibitors of host translation from *S. enterica* phages follow the same trend (Figure 5c). Interestingly, small proteins such as the inhibitors of the host toxin/antitoxin system from *S. enterica* and the anti-sigma factors from *E. coli* phages have unique structures (Figure 5d). This protein structure analysis of the representative proteins with potential antimicrobial activities reveals the underexplored surface of the protein structure landscape, which are small proteins with diverse structural components (Figure 1a).

## Discussion

The underrepresentation of bacteriophages in biological databases despite being the most abundant biological entities in the biosphere raises concerns that a significant portion of this vital biosphere remains unexplored. For example, the protein structure landscape is not complete without exploring the small proteins from phages (Figure 1a). These small proteins derived from phages are increasingly recognized for their remarkable diversity [12,41]. This study seeks to quantify the extent of the underrepresentation of phage data within public databases and to discuss the implications of such biased datasets. Specifically, it aims to assess the impact on medically relevant fields such as phage therapy, as well as on data-driven computational models like deep learning-based protein structure prediction programs.

In this study, we revealed the extent of underrepresentation of bacteriophages in the Reference Sequence (RefSeq) of the NCBI Virus. This database provides a comprehensive and non-redundant set of sequences and a stable reference for genome annotation, forming a foundation for medical and functional studies. We examined the phage genomes infecting 11 bacterial species categorized under the ESKAPE acronym, designated as the top priority pathogens by the WHO for new drug development due to widespread multidrug resistance [8]. The gene content analysis shows that several ESKAPE pathogens, such as *C. jejuni*, *E. faecium*, and *H. influenzae*, have only a few phages that are likely to have antimicrobial activities. Notably, *H. pylori* has no unique phages with functionally annotated genes encoding for antimicrobial activities (Figure S1). More importantly, these bacteria have no putative antimicrobial genes (Figure S2).

Next, we conducted the lifestyle analysis of these ESKAPE phages to confirm that there is only a handful of lytic phages for most pathogens (Table S2). The lifestyle of bacteriophages holds profound implications across various fields, including phage therapy, genomics, and microbiology. In the lytic lifecycle, the phage infects a bacterial host cell, hijacks its machinery to replicate its own genetic material and produce progeny phages, and ultimately causes the host cell to burst, releasing the newly formed phages to infect neighboring cells. This process results in the immediate destruction of the host cell [3]. On the other hand, in the lysogenic lifecycle, the phage inserts its genetic material into the host cell’s genome, becoming a prophage. The prophage is replicated along with the host cell’s DNA during cell division, remaining latent within the host cell without causing immediate harm. Under certain conditions, such as exposure to stressors, the prophage may become activated, entering the lytic cycle and causing the host cell to lyse [54]. The lack of lytic phages in the genomic repertoire reduces the number of treatment options in phage therapy [8]. Phage therapy, leveraging the natural predation of bacteriophages on bacteria, is gaining traction as a viable alternative or complementary strategy to traditional antibiotics [6,7]. As antibiotics struggle to maintain efficacy against evolving bacterial defenses, bacteriophages offer a tailored and evolving solution, capable of adapting to bacterial mutations [8].

Given the lack of phages with antimicrobial activities, we used the pangenome analysis to reveal how many of the ESKAPE phages are unique in terms of nucleotide identity and gene content. From whole-genome clustering and average nucleotide identity computation, we found that the ESKAPE phages of the same host share many core genes. Additionally, the ESKAPE phages form a small number of clusters in the whole-genome phylogenetic analysis. The average nucleotide identity analysis also highlights a deficiency in unique phage genomes within the reference sequence database, which ideally should offer a comprehensive and non-redundant collection of sequences. To enhance the classification of phage genomes, we propose the adoption of clustering methods that prioritize gene content rather than gene arrangement, considering the rapid evolutionary rate of phages.

Last, we used the deep learning-based structure program to predict the three-dimensional structures of the representative proteins with potential antimicrobial activities in the ESKAPE phages. After clustering with the multiple sequence alignment method, we selected representative sequences for each cluster (Figure S6). From the visualization, we found that the ESKAPE phages only possess a few antimicrobial proteins with unique structures (Figure 5). Notably, the unique structures of these antimicrobial proteins are mostly small proteins such as anti-sigma proteins or toxin/antitoxin proteins.

The sequencing of bacteriophages presents distinct challenges compared to other microbial entities, such as bacteria and archaea. Bacteriophages exhibit an extraordinary degree of genetic diversity. Unlike bacteria and archaea, for which reference genomes are relatively abundant, the lack of comprehensive reference databases for bacteriophages complicates the alignment and assembly processes during sequencing [55]. Bacteriophages exhibit rapid rates of evolution, leading to genomic changes that occur over short periods [56]. This evolutionary dynamism can complicate the assembly and analysis of phage genomes, particularly when attempting to capture the full spectrum of genetic variation. Environmental samples often contain a multitude of different bacteriophages, and some phages can even infect the same bacterial host. This phenomenon, known as mixed infections, introduces complexities in the interpretation of sequencing data, making it challenging to distinguish individual phage genomes in a mixture [57]. Other challenges arise from a combination of factors, including limitations in phage-specific extraction protocols and computational tools. Addressing these challenges in bacteriophage sequencing requires the development of specialized protocols, bioinformatics tools, and reference databases tailored to the unique characteristics of these viral entities. Recent advances in sequencing technologies and methodologies are continually improving our ability to overcome these obstacles and unravel the genetic intricacies of bacteriophages [58].

Lastly, we emphasize the implications of biased datasets on data-driven computational models, particularly deep learning-based tools that have been trained on the currently available datasets for decision-making and generative processes [12,41]. The challenge of underrepresented datasets poses a substantial concern, particularly in the context of machine learning models. This issue becomes particularly pronounced when the training data used to train these models is biased, leading to skewed and potentially inaccurate results during the model’s predictive or classification tasks. In machine learning, the efficacy and reliability of a model are highly contingent on the quality and representativeness of the data it is trained on. When certain groups or categories within the dataset are underrepresented, the model may not adequately learn the patterns and characteristics associated with those groups [9,10]. This lack of representation can lead to a biased understanding of the data, resulting in a model that is less capable of making accurate predictions or classifications for the underrepresented groups. The consequences of biased training data are far-reaching and can manifest in various ways. For instance, in predictive modeling of protein structures, the model may struggle to generalize well to instances that belong to underrepresented classes, leading to poor performance for phage proteins. Bacteriophages have undergone a paradigm shift in recent years. Initially perceived as having limited medical relevance compared to bacteria, bacteriophages are now recognized as essential players in various medical and therapeutic contexts. This shift in perspective is primarily attributed to a deeper understanding of the intricate relationships between bacteriophages and their bacterial hosts.

## Declarations

### Availability of data and materials

All codes related to this project are available under an open-source license at https://github.com/hshimlab. For data analysis, Python v.3.6.4 (https://www.python.org), NumPy v.1.17.5 (https://github.com/numpy/numpy), SciPy v.1.1.0 (https://www.scipy.org), seaborn v.0.9.0 (https://github.com/mwaskom/seaborn), Matplotlib v.3.3.4 (https://github.com/matplotlib/matplotlib), pandas v.0.22.0 (https://github.com/pandas-dev/pandas) were used. Genome analyses were conducted with PHACTS (http://www.phantome.org/PHACTS), MMseqs2 (https://github.com/soedinglab/MMseqs2), and IPGA v1.09 (https://nmdc.cn/ipga). Protein structures were predicted with AlphaFold2, available under an open-source license at https://github.com/deepmind/alphafold. As protein structure similarity metrics, we used TM-align (https://zhanggroup.org/TM-align). 3-D structure visualizations were created with 3Dmol.js (https://3dmol.csb.pitt.edu/doc/tutorial-embeddable.html).

## References

1. Suttle CA. Viruses in the sea. Nature. 2005;437: 356–361.

2. Clokie MRJ, Millard AD, Letarov AV, Heaphy S. Phages in nature. Bacteriophage. 2011;1: 31.

3. Shim H, Shivram H, Lei S, Doudna JA, Banfield JF. Diverse ATPase Proteins in Mobilomes Constitute a Large Potential Sink for Prokaryotic Host ATP. Front Microbiol. 2021;12: 691847.

4. Al-Shayeb B, Sachdeva R, Chen L-X, Ward F, Munk P, Devoto A, et al. Clades of huge phages from across Earth’s ecosystems. Nature. 2020;578: 425–431.

5. Global burden of bacterial antimicrobial resistance in 2019: a systematic analysis. Lancet. 2022;399: 629–655.

6. Gordillo Altamirano FL, Barr JJ. Phage Therapy in the Postantibiotic Era. Clin Microbiol Rev. 2019;32. doi:10.1128/CMR.00066-18

7. Young R, Gill JJ. Phage therapy redux—What is to be done? Science. 2015 [cited 14 Feb 2024]. doi:10.1126/science.aad6791

8. Shim H. Three Innovations of Next-Generation Antibiotics: Evolvability, Specificity, and Non-Immunogenicity. Antibiotics (Basel). 2023;12. doi:10.3390/antibiotics12020204

9. Deviyani A. Assessing Dataset Bias in Computer Vision. 2022 [cited 18 Feb 2024]. doi:10.13140/RG.2.2.19950.89924

10. Jones C, Castro DC, De Sousa Ribeiro F, Oktay O, McCradden M, Glocker B. A causal perspective on dataset bias in machine learning for medical imaging. Nature Machine Intelligence. 2024; 1–9.

11. Dill KA, Ozkan SB, Shell MS, Weikl TR. The protein folding problem. Annu Rev Biophys. 2008;37. doi:10.1146/annurev.biophys.37.092707.153558

12. Jumper J, Evans R, Pritzel A, Green T, Figurnov M, Ronneberger O, et al. Highly accurate protein structure prediction with AlphaFold. Nature. 2021;596: 583–589.

13. Santajit S, Indrawattana N. Mechanisms of Antimicrobial Resistance in ESKAPE Pathogens. Biomed Res Int. 2016;2016. doi:10.1155/2016/2475067

14. Berman HM, Westbrook J, Feng Z, Gilliland G, Bhat TN, Weissig H, et al. The Protein Data Bank. Nucleic Acids Res. 2000;28: 235–242.

15. ChatGPT: A comprehensive review on background, applications, key challenges, bias, ethics, limitations and future scope. Internet of Things and Cyber-Physical Systems. 2023;3: 121–154.

16. Prioritization of pathogens to guide discovery, research and development of new antibiotics for drug-resistant bacterial infections, including tuberculosis. World Health Organization; 2019.

17. Galperin MY. Conserved “Hypothetical” Proteins: New Hints and New Puzzles. Comp Funct Genomics. 2001;2: 14.

18. Alexandre S, Guyaux M, Murphy NB, Coquelet H, Pays A, Steinert M, et al. Putative genes of a variant-specific antigen gene transcription unit in Trypanosoma brucei. Mol Cell Biol. 1988;8: 2367.

19. McNair K, Bailey BA, Edwards RA. PHACTS, a computational approach to classifying the lifestyle of phages. Bioinformatics. 2012;28. doi:10.1093/bioinformatics/bts014

20. Ho TK. Random decision forests. [cited 1 Feb 2024]. Available: https://ieeexplore.ieee.org/abstract/document/598994

21. Ho TK. The random subspace method for constructing decision forests. [cited 1 Feb 2024]. Available: https://ieeexplore.ieee.org/abstract/document/709601

22. Batstone RT, Burghardt LT, Heath KD. Phenotypic and genomic signatures of interspecies cooperation and conflict in naturally occurring isolates of a model plant symbiont. Proc Biol Sci. 2022;289: 20220477.

23. Liu D, Zhang Y, Fan G, Sun D, Zhang X, Yu Z, et al. IPGA: A handy integrated prokaryotes genome and pan-genome analysis web service. iMeta. 2022;1: e55.

24. Zhou Z, Charlesworth J, Achtman M. Accurate reconstruction of bacterial pan- and core genomes with PEPPAN. Genome Res. 2020;30: 1667.

25. Konstantinidis KT, Tiedje JM. Genomic insights that advance the species definition for prokaryotes. Proc Natl Acad Sci U S A. 2005;102. doi:10.1073/pnas.0409727102

26. Steinegger M, Söding J. MMseqs2 enables sensitive protein sequence searching for the analysis of massive data sets. Nat Biotechnol. 2017;35: 1026–1028.

27. Fleischmann RD, Adams MD, White O, Clayton RA, Kirkness EF, Kerlavage AR, et al. Whole-Genome Random Sequencing and Assembly of Haemophilus influenzae Rd. Science. 1995 [cited 17 Feb 2024]. doi:10.1126/science.7542800

28. Michniewski S, Rihtman B, Cook R, Jones MA, Wilson WH, Scanlan DJ, et al. A new family of “megaphages” abundant in the marine environment. ISME Communications. 2021;1: 1–4.

29. Fremin BJ, Bhatt AS, Kyrpides NC, Global Phage Small Open Reading Frame (GP-SmORF) Consortium. Thousands of small, novel genes predicted in global phage genomes. Cell Rep. 2022;39: 110984.

30. Silpe JE, Duddy OP, Johnson GE, Beggs GA, Hussain FA, Forsberg KJ, et al. Small protein modules dictate prophage fates during polylysogeny. Nature. 2023;620: 625–633.

31. Zhang S, Gu J, Shi L-L, Qian B, Diao X, Jiang X, et al. A pan-cancer analysis of anti-proliferative protein family genes for therapeutic targets in cancer. Sci Rep. 2023;13: 1–15.

32. Kalie E, Jaitin DA, Abramovich R, Schreiber G. An Interferon α2 Mutant Optimized by Phage Display for IFNAR1 Binding Confers Specifically Enhanced Antitumor Activities *. J Biol Chem. 2007;282: 11602–11611.

33. Paget MS. Bacterial Sigma Factors and Anti-Sigma Factors: Structure, Function and Distribution. Biomolecules. 2015;5: 1245.

34. Pilotto S, Fouqueau T, Lukoyanova N, Sheppard C, Lucas-Staat S, Díaz-Santín LM, et al. Structural basis of RNA polymerase inhibition by viral and host factors. Nat Commun. 2021;12: 1–15.

35. Merrikh H, Zhang Y, Grossman AD, Wang JD. Replication-transcription conflicts in bacteria. Nat Rev Microbiol. 10: 449.

36. Spoerel N, Herrlich P, Bickle TA. A novel bacteriophage defence mechanism: the anti-restriction protein. Nature. 1979;278: 30–34.

37. Meyn MS, Rossman T, Troll W. A protease inhibitor blocks SOS functions in Escherichia coli: antipain prevents lambda repressor inactivation, ultraviolet mutagenesis, and filamentous growth. Proc Natl Acad Sci U S A. 1977;74. doi:10.1073/pnas.74.3.1152

38. Horiuchi T, Sakamoto M, Murotsu T. Studies on lambda virulent mutants. III. Action of the anti- and vir-repressor (cro-product) of lambda phage on the related lambdoid phages. Mol Gen Genet. 1974;133. doi:10.1007/BF00268677

39. Bondy-Denomy J, Pawluk A, Maxwell KL, Davidson AR. Bacteriophage genes that inactivate the CRISPR/Cas bacterial immune system. Nature. 2012;493: 429–432.

40. Park H-M, Park Y, Berani U, Bang E, Vankerschaver J, Van Messem A, et al. In silico optimization of RNA-protein interactions for CRISPR-Cas13-based antimicrobials. Biol Direct. 2022;17: 27.

41. Park H-M, Park Y, Vankerschaver J, Van Messem A, De Neve W, Shim H. Rethinking Protein Drug Design with Highly Accurate Structure Prediction of Anti-CRISPR Proteins. Pharmaceuticals. 2022;15: 310.

42. Shim H. Investigating the genomic background of CRISPR-Cas genomes for CRISPR-based antimicrobials. arXiv [q-bio.GN]. 2022. Available: http://arxiv.org/abs/2202.07171

43. Li L, Stoeckert CJ Jr, Roos DS. OrthoMCL: identification of ortholog groups for eukaryotic genomes. Genome Res. 2003;13: 2178–2189.

44. Santos AR, Barbosa E, Fiaux K, Zurita-Turk M, Chaitankar V, Kamapantula B, et al. PANNOTATOR: an automated tool for annotation of pan-genomes. Genet Mol Res. 2013;12: 2982–2989.

45. Page AJ, Cummins CA, Hunt M, Wong VK, Reuter S, Holden MTG, et al. Roary: rapid large-scale prokaryote pan genome analysis. Bioinformatics. 2015;31: 3691–3693.

46. Fouts DE, Brinkac L, Beck E, Inman J, Sutton G. PanOCT: automated clustering of orthologs using conserved gene neighborhood for pan-genomic analysis of bacterial strains and closely related species. Nucleic Acids Res. 2012;40: e172.

47. DeSalle R, Tessler M, Rosenfeld J. Phylogenomics: A Primer. CRC Press; 2020.

48. Chen X, Zhang Y, Zhang Z, Zhao Y, Sun C, Yang M, et al. PGAweb: A Web Server for Bacterial Pan-Genome Analysis. Front Microbiol. 2018;9: 1910.

49. Ding W, Baumdicker F, Neher RA. panX: pan-genome analysis and exploration. Nucleic Acids Res. 2018;46: e5.

50. Gautreau G, Bazin A, Gachet M, Planel R, Burlot L, Dubois M, et al. PPanGGOLiN: Depicting microbial diversity via a partitioned pangenome graph. PLoS Comput Biol. 2020;16: e1007732.

51. Tonkin-Hill G, MacAlasdair N, Ruis C, Weimann A, Horesh G, Lees JA, et al. Producing polished prokaryotic pangenomes with the Panaroo pipeline. Genome Biol. 2020;21: 180.

52. Emms DM, Kelly S. OrthoFinder: phylogenetic orthology inference for comparative genomics. Genome Biol. 2019;20: 238.

53. Galperin MY, Wolf YI, Makarova KS, Vera Alvarez R, Landsman D, Koonin EV. COG database update: focus on microbial diversity, model organisms, and widespread pathogens. Nucleic Acids Res. 2021;49: D274–D281.

54. Knowles B, Silveira CB, Bailey BA, Barott K, Cantu VA, Cobián-Güemes AG, et al. Lytic to temperate switching of viral communities. Nature. 2016;531: 466–470.

55. Shim H. Futuristic Methods in Virus Genome Evolution Using the Third-Generation DNA Sequencing and Artificial Neural Networks. Global Virology III: Virology in the 21st Century. 2019; 485–513.

56. Shim H. Feature Learning of Virus Genome Evolution With the Nucleotide Skip-Gram Neural Network. Evol Bioinform Online. 2019 [cited 19 Feb 2024]. doi:10.1177/1176934318821072

57. Mathew S, Smatti MK, Al Ansari K, Nasrallah GK, Al Thani AA, Yassine HM. Mixed Viral-Bacterial Infections and Their Effects on Gut Microbiota and Clinical Illnesses in Children. Sci Rep. 2019;9: 1–12.

58. Park Y, Lee J, Shim H. Sequencing, Fast and Slow: Profiling Microbiomes in Human Samples with Nanopore Sequencing. Applied Biosciences. 2023;2: 437–458.

